# Covalent on-cell conjugation of biomaterials through oxidative phenolic coupling regulates stem cell fate via intracellular biophysical programming

**DOI:** 10.1101/2024.09.23.614446

**Authors:** Castro Johnbosco, Malin Becker, Niels Willemen, Marieke Meteling, Kannan Govindaraj, Tom Kamperman, Jeroen Leijten

## Abstract

Mechanotransduction is widely used to guide cell fate in hydrogels. Traditionally, hydrogels contain adhesive ligands that dynamically bond with cells to stimulate biochemical signalling axis such as YAP-TAZ. However, the molecular toolbox to achieve mechanotransduction has remained virtually limited to non-covalent bonds, which limits our ability to program engineered living matter. Here, we demonstrate that on-cell chemistry can be leveraged to covalently tether biomaterials directly onto cells, and reveal that mechanotransduction is enabled via intracellular biophysical programming. Specifically, droplet microfluidics produced single-cell microgels in which individual stem cells were extracellularly conjugated to either soft or stiff hydrogels via on-cell oxidative phenolic coupling, which allowed for investigation of mechanotransduction at single-cell resolution. Interestingly, this altered intracellular molecular crowding, calcium signalling, and chromatin organization by regulating cytoplasmic and nuclear volume in a stiffness-dependent yet YAP/TAZ-independent manner. Notably, addition of conventional dynamic adhesive ligands such as RGDs decreased chondrogenic commitment indicating that covalent cell-material tethering is both efficient and sufficient for programming cell fate. Encoding biomaterials with a novel form of mechanotransduction in the form of covalent on-cell chemistry, such as oxidative phenolic coupling, expands our ability to guide cellular behaviour, which can accelerate development of drug-screening models, lab-grown meat, and engineered tissues.

## Introduction

The mechanical properties of the pericellular matrix determine various cellular processes including self-renewal^1–3^, cell differentiation^4–6^, pathological onset, and disease progression^7,8^. Engineering living tissues composed of cells in biomaterials is of high importance for a variety of applications including basic science research^9^, production of transplantable living organs^10^, development of drug screening models^11^, and biofabrication of lab-grown meat^12^. Most commonly, cells are placed in a matrix in which mechanotransduction is mediated via cell-adhesive moieties to guide cell fate and control tissue function^13^. However, the material toolbox to endow materials with cell adhesive properties has remained limited to a few distinct types of molecules, all of which form non-covalent, reversible, and dynamic bonds^14^. For example, the tripeptide Arg-Gly-ASP (RGD) is the most commonly investigated cell-binding moiety, which facilitates cellular-material interactions by weakly and reversibly bonding integrins^15,16^. Other examples of explored cell-binding moieties include distinct peptides, nucleotides^17^, aptamers^18^, and antibodies^19^. However, despite being an effective manner to interact with a limited set of cellular ligands and their associated intracellular signaling cascades, this approach poorly mimics the natural abundance of variety in cell-materials interactions experienced by cells in their native microenvironments^20^.

On-cell chemistry has recently been introduced as a novel method to achieve cell-material interactions typically by offering more stable and even covalent cell-material bonds^21,22^. Advantageously, cell surfaces are decorated with binding sites^23^ with various functional groups ^24,25^ that offer space for covalent on-cell conjugation of materials. Cell surface engineering approaches could thus be leveraged to establish engineered covalent, non-dynamic mechanotransduction routes^26^, which offer attractive properties to study and guide (stem) cell fate. To achieve such precise covalent tether-mediated mechanotransduction, we explored oxidative phenolic crosslinking to achieve discrete inducible on-cell crosslinking (DOCKING)^27^. This approach effectively tethers tyramine-conjugated to the tyrosine that cells naturally present in their natural pericellular matrix and transmembrane proteins^28^. However, how this novel form of mechanotransduction governs changes in cell function and fate changes has remained largely unknown.

Here, we report that on-cell covalent bonding of extracellular matrix via oxidative phenolic crosslinking biophysically alters the intracellular organization of stem cells, which guides the programming of stem cell fate. To gain these insights at single cell level in 3D microenvironments, we leveraged a microfluidic chip-based platform to produce single-cell microgels, which were composed of individual cells that were uniformly and covalently coated with a thin layer of hydrogel (∼5µm). The stiffness of this micrometer thin layer of the hydrogel could be on-demand tuned, which was used to investigate the optimal stiffness to guide chondrogenic lineage commitment of bone marrow-derived stem cells. This demonstrated that oxidative phenolic crosslinking-mediated covalent mechanotransduction in chondrogenic induction was correlated with changes in intracellular molecular crowding, calcium signaling, and the nucleus’ biophysical, protein, and histone arrangements. Interestingly, unlike conventional dynamic non-covalent mechanosensing strategies, oxidative phenolic crosslinking-mediated proved independent of YAP/TAZ nuclear translocation. Moreover, potential complementary effects addition of conventional non-covalent dynamic binder (e.g., RGDs) were investigated, which revealed that the introduction of non-covalent binders significant into covalent oxidative phenolic crosslinking notably inhibited the chondrogenic commitment of stem cells. Taken together, this study reveals that the covalent and static mechanotransduction route operates over distinct communication axis from known non-covalent binder-induced mechanotransduction mechanisms, which operate over biophysical cellular alterations to alter the lineage commitment of stem cells.

## Results

### Microfluidic fabrication of single-cell microgels with tunable stiffness using enzyme-mediated oxidative phenolic crosslinking

To engineer customizable pericellular microenvironments for individual single cells, we leveraged an in-house designed droplet-based microfluidic platform operated in flow focus mode in which a continuous oil phase and discrete aqueous phase are co-flown to form a water-in-oil emulsion^29^. To this end, Dextran was chosen owing to it being a biocompatible polymer commonly used to study cell-material interactions. Dextran was functionalized with tyramine (DexTA; Ds 14%) (Figure S1) to allow for mechanotransduction via DOCKING achieved through enzyme-mediated oxidative phenolic crosslinking, which is catalyzed by horseradish peroxidase (HRP) in the presence of the oxidizing hydrogen peroxide. Briefly, DexTA solution containing human bone marrow-derived mesenchymal stromal cells (MSCs) was emulsified into droplets and subsequently outside-in crosslinked to form conformally coated single-cell microgels. Outside-in crosslinking of polymer microdroplets was achieved by diffusing hydrogen peroxides via parallel channels through PDMS walls and the emulsion’s oil phase. This process allows for highly controlled dosing of hydrogen peroxide into the discrete aqueous droplets thereby inducing tyramine crosslinking via covalent C-C and C-O bond formation (Figure 1a). By altering the volume ratios of dispersed aqueous flow and continuous oil phase as a function of time, we could obtain accurate control of droplet volume in a predictable and monodisperse manner (Figure 1b-d). Controlled supplementation of hydrogen peroxide also enabled accurate control over crosslink density and thus elastic modulus, which we here leveraged to produce ∼14 kPa (soft) and ∼38 kPa (stiff) single-cell microgels (Figure 1e). Dynamic mechanical analysis at various frequencies revealed a stable storage modulus with a mild increasing trend among both soft and stiff microgels (Figure 1f). EthD-1 staining of oxidative phenolic crosslinked microgels is known to directly correlate with crosslink density^27^, which corroborates that the difference between soft and stiff microgels is associated with differences in tyramine dimer density (Figure 1g). To investigate intra-microgel spatial variation in crosslink density, mechanical properties were semi-quantified in a spatially resolved manner, which revealed near-neglectable spatial variations in crosslink densities for both soft (CV 10.7%) and stiff (CV 5.8%) microgels (Figure 1h). Of note, the EthD-1 fluorescent signal exhibited a strong linear correlation (R² = 0.99) with the microgel’s Young modulus, which corroborated its relation with crosslink density (Figure S2). To confirm that the altered crosslink density did not majorly affect the diffusion rates, microgels were submerged in 20 kDa FITC-Dextran for up to 24 hours (Figure 1i). Although a minor stiffness-dependent effect was observed, fluorophores freely diffused throughout both soft and stiff microgels within a few hours (Figure 1j). This suggests that nutrients, waste products, and cytokines can freely diffuse through the microgels regardless of their stiffness. In short, this approach offers controlled microfluidic production of mechanically tunable microgel platforms to enable the study of cellular biological processes in customizable 3D microenvironments at single-cell resolution.

**Figure 1:**
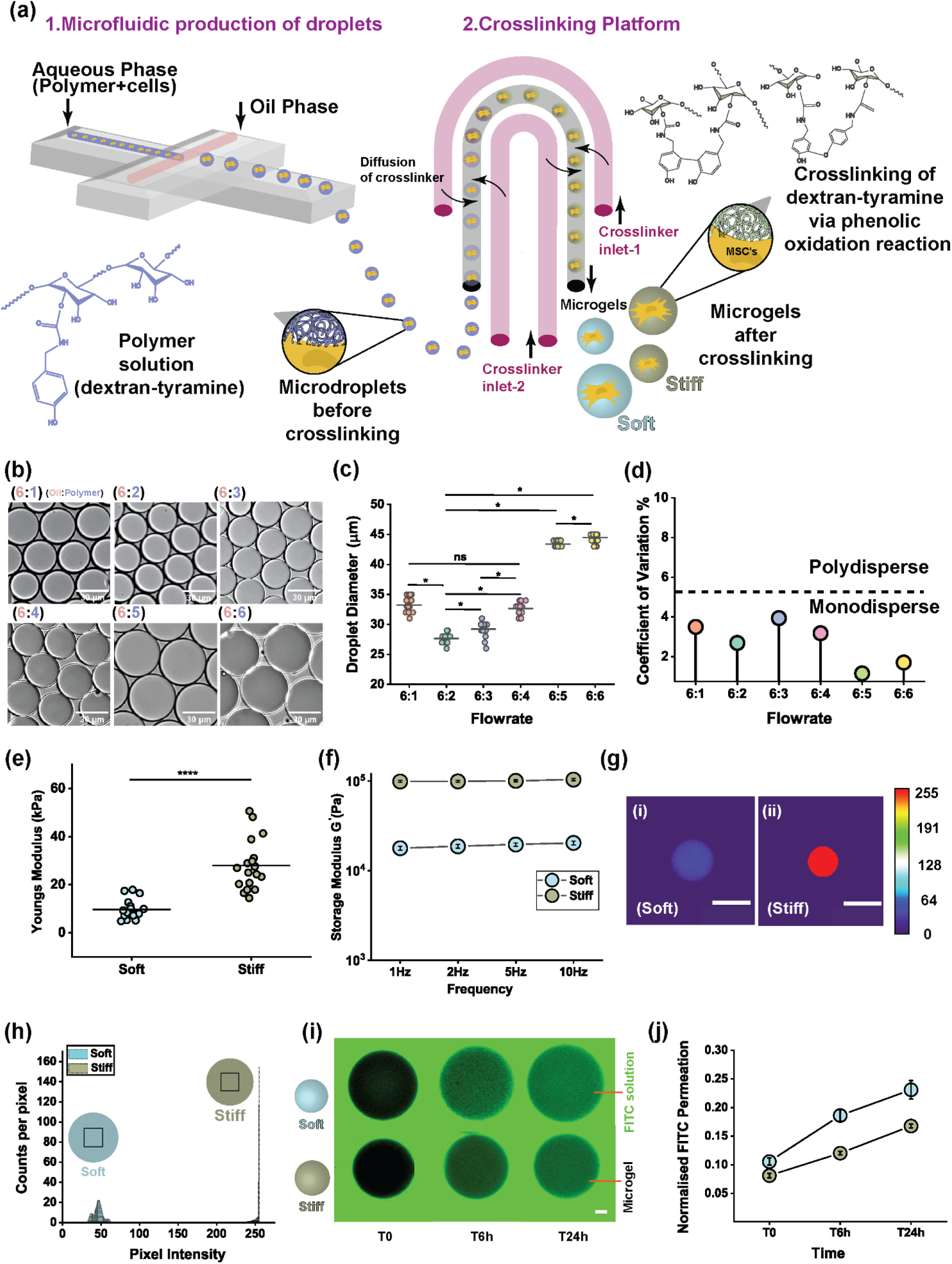
Microfluidic production of mechanically tunable microgels. (**a**) Schematic depiction of microdroplets and (single cell) microgel production using an in-line dual microfluidic chip platform based on a flow-focus microfluidic droplet generator and outside-in enzymatic oxidative phenolic crosslinking. (**b**) Brightfield micrographs of droplets, (**c**) image-based quantification of droplet diameter (n=20), and (**d**) coefficient of variation of diameter droplets, produced at different flow rates (oil, red; polymer solution, white) (n=20). (Scale bar indicates 30 µm). *p<0.05 by one-way ANOVA (**e**) Youngs modulus of microgels was determined using Interferometry-based mechanical nanoindentation analysis (n=19). ****p<0.0001 by one-way ANOVA. (**f**) DMA analysis on soft and stiff microgels at varying frequencies ranging from 1 Hz to 10 Hz. (**g**) Total fluorescence intensity quantification of EthD-1 stained (i) soft or (ii) stiff microgels (a.u). (**h**) Per pixel image analysis of EthD-1 stained microgels to semi-quantify intra-microgel stiffness variation within soft and stiff microgels (n≥15). (**i**) Fluorescent micrographs of microgels submerged in 20 kDa FITC-Dextran (green) for up to 24 hours (Scale bar indicates 5 µm). (**j**) Normalized FITC intensity quantification inside soft and stiff microgels (a.u)(n≥17). All data shown are mean ± s.e.m.

### Tethering biomaterial tyramine conjugates to extracellular tyrosines enables mechanotransduction

For engineered microniches to mechanically instruct cells, it is essential to enable mechanotransduction to facilitate this cell-material interaction. Various reversible, non-covalent, continuously active, adhesion-dependent strategies to allow cells to sense their microenvironmental stiffness have been investigated^14,30^. In contrast to previous approaches, enzymatic oxidative phenolic crosslinking allows for the on-cell formation of inducible covalent cell-material bonds. This was achieved via the coupling of biomaterials decorated with tyramines to tyrosines, which are naturally available amino acids expressed in transmembrane proteins and secreted extracellular matrix proteins of (mammalian) cells (Figure 2a). Tyrosine is chemically an analog of tyramine but exists in a carboxylated state, hence tyrosine will form phenolic crosslinks with tyramine when in the presence of HRP and H_2_O_2_ (Figure 2b). To validate this concept, an equal molar mixture of tyramine and tyrosine was crosslinked in the presence of HRP and H_2_O_2_, while H_2_O was used instead of H_2_O_2_ for the negative control. UV-vis spectroscopy (Figure 2c) revealed that in the presence of H_2_O_2_ tyramine and tyrosine exhibited a hyperchromic shift at 221 nm, a peak disappearance at 241nm, and a peak appearance at 314 nm, which did not occur in the presence of H_2_O. Together this demonstrated that tyramine-tyrosine molecules were crosslinked via phenolic oxidation in the presence of HRP and H_2_O_2_ thus being able to effectively covalently tether tyramine-modified materials directly onto cells to facilitate mechanotransduction.

**Figure 2:**
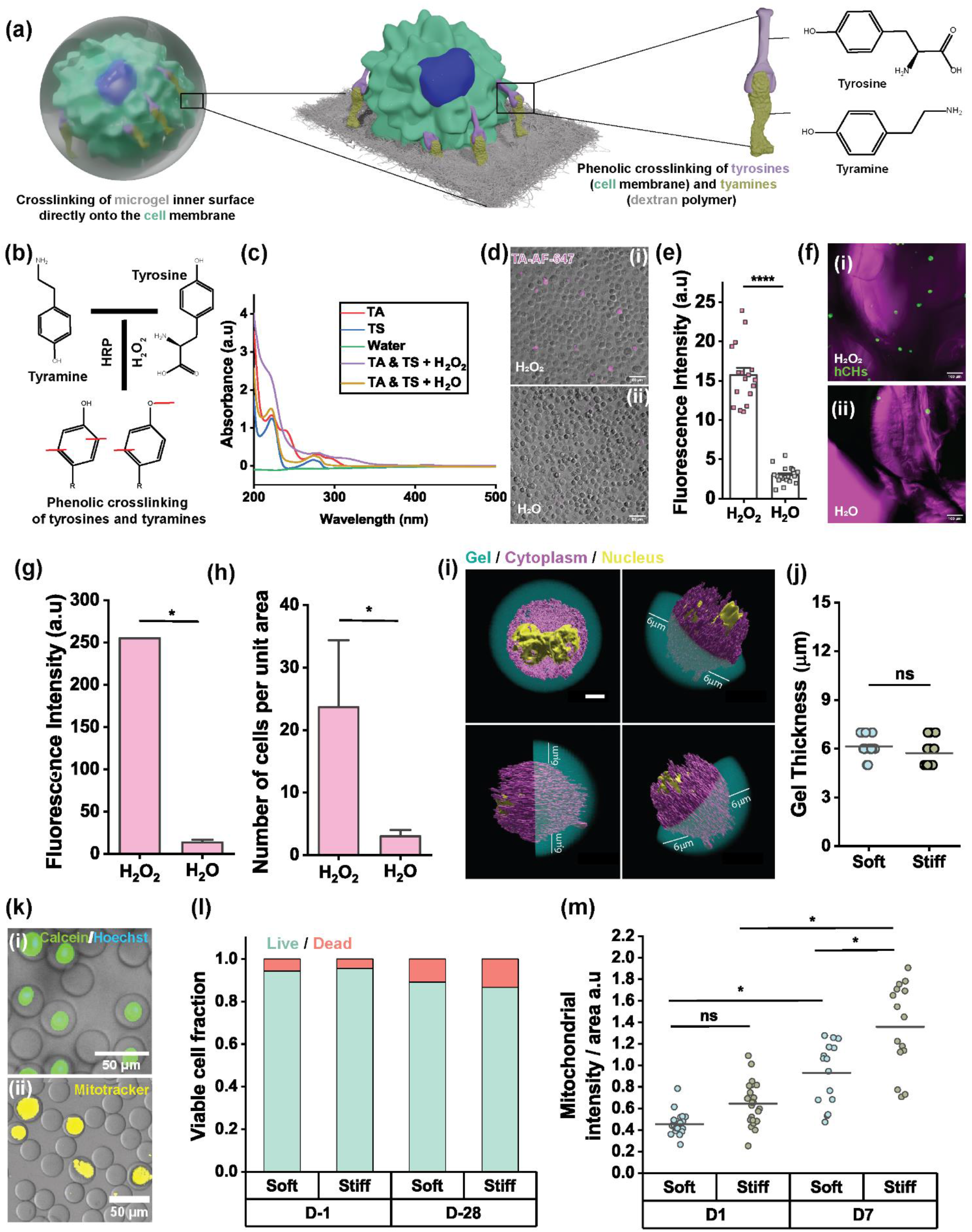
Discrete inducible on-cell crosslinking via oxidative phenolic crosslinks tethers cells to microgels. **(a)** Schematic depiction of dextran-tyramine polymer conjugates tethering to extracellular tyrosines. **(b)** Schematic depiction and **(c)** UV-vis spectra of phenolic crosslinking of tyramine-tyrosine molecules catalyzed by HRP in the presence of H_2_O_2_. **(d)** Fluorescence micrographs of TA AF-647 coupled to cells in the presence of (i) H_2_O_2_, (ii) H_2_O, and **(e)** quantitation of fluorescence signal intensity of human chondrocytes (hCHs) exposed to AF-647 tyramide in the presence of either H_2_O_2_ or H_2_O to demonstrate covalent enzymatic phenolic tethering. (****p<0.0001 by Kruskal-Wallis ANOVA) (n ≥ 17). **(f)** Confocal micrographs of hCHs (green) tethered to DexTA hydrogel discs in the presence of (i)H_2_O_2_ and (ii) H_2_O (n=3 per condition) **(g)** Total fluorescence intensity calculation on dextran-tyramine hydrogels, **(h)** number cells tethered to dextran-tyramine hydrogels (n=3 per condition). **(i)** 3D reconstruction of confocal fluorescence micrographs of single cell microgels including the interior of the microniches, orthographic, and isometric confirming equidistant positioning of MSCs from the microgel periphery (Scale bar indicates 4 µm). **(j)** Confocal image quantified microgel shell thickness for both soft and stiff single cell microgels (n=15). **(k)** Fluorescence micrographs of single-cell microgels stained with (i) calcein (green), Hoechst (blue), and (ii) mitotracker (yellow) to determine cell survival and mitochondrial concentration. **(l)** Image-quantified fraction of viable cells in soft and stiff single-cell microgels (n≥40). **(m)** Quantified mitochondrial concentration per single cell microgel to determine mitochondrial activity (n≥15) (*p<0.05 by Kruskal-Wallis ANOVA). All data shown are mean± s.e.m

To demonstrate the oxidative phenolic coupling of tyramines to cells, a mixture of fluorophore tyramide conjugates (TA-AF-647) and HRP was exposed to cells in the presence of either H_2_O_2_ or H_2_O. As anticipated, cells were covalently and stably painted with the fluorophore in the presence of H_2_O_2_ but remained pristine when exposed instead to H_2_O (Figure 2d & 2e). Moreover, exposing cells atop on Dex-TA hydrogel disc to H_2_O_2_ resulted in the stable tethering of cells to the hydrogel surface (Figure 2f). This allowed tethered cells to withstand mild hydrodynamic agitation while non-tethered cells (e.g., those exposed to H_2_O) readily washed off from the hydrogel surface (Figure 2g & 2h). Together, this corroborated that cells can be enzymatically tethered to materials via phenolic oxidative crosslinks by forming tyrosine-tyramine bonds. To translate this strategy towards a 3D culture platform that offers inherent single-cell resolution analyses, microfluidic droplet generators were used to encapsulate individual MSCs in 30 µm microgels, which could be controllably produced to offer either soft or stiff microenvironments. The use of a previously reported modular delayed crosslinking platform^31^ produced single-cell microgels that conformally encapsulated each cell as evidenced by 3D reconstructed images, which demonstrated equidistant positioning in both orthographic and isometric views (Figure 2i). Consequently, all single-cell microgels possessed a uniform microgel shell thickness, which was independent of microgel stiffness (Figure 2j). Microencapsulation of MSCs associated with a high survival rate and maintenance of metabolic activity as demonstrated via continued mitochondrial activity in both soft and stiff microgels for at least 28 days of culture (Figure 2k). Specifically, upon microencapsulation >95% of MSCs survived, and ∼90% of MSCs were still viable after 28 days of culture, which was not affected by the use of either soft or stiff microgels (Figure 2l). Moreover, the cytocompatibility of the approach was further corroborated by the observation that mitochondrial activity nearly doubled within seven days of culture, regardless of microgel stiffness (Figure 2m). This suggests that mechanosensing of the surrounding pericellular stiffness by MSCs inside microgels mediated via oxidative phenolic crosslinking is sufficient to alter cellular performance including metabolism by influencing amongst others mitochondrial activity.

### Soft pericellular environments promote chondrogenic lineage commitment via phenolic coupling

We next investigated whether covalent tethering of cells to their physical microenvironment via oxidative phenolic crosslinking could orchestrate lineage commitment of microencapsulated stem cells. To this end, MSCs were encapsulated in either soft or stiff microgels and exposed to a chondrogenic differentiation medium. To evaluate differences in nascent protein deposition in single-cell microgels, we innovatively explored the use of holotomography imaging, which is a novel label-free method based on refractive index quantitation using quantitative phase imaging to determine the amount of protein produced in a high-resolution and three-dimensional spatially resolved manner^32,33^. After two weeks of chondrogenic differentiation, MSCs inside soft microgels surprisingly had deposited notably higher levels of nascent proteins than MSCs inside stiff microgels (Figure 3a). Holotomography revealed that the refractive index of MSCs in stiff microgels was not noticeably decreased, while that of MSCs in soft microgels had significantly increased (Figure 3b). Moreover, the protein concentration per single-cell microgel was numerically computed based on the refractive index value ^32^ for various time points, revealing the deposition of ∼5 g/dl protein for MSCs in soft microgels (Figure 3c & 3d). To corroborate these label-free findings, single-cell microgels were also investigated using immunohistochemistry and immunofluorescence confocal imaging of specific matrix proteins related to chondrogenic differentiation including COL2, ACAN, COL6, and SOX9 (Figure 3e). Single stem cells in soft microgels had significantly higher fluorescence intensity for cartilage’s major matrix components COL2 (Figure 3f) and ACAN (Figure 3g) compared to cells in stiff microgels. COL6, which is a pericellular matrix protein expressed by chondrocytes, showed no significant difference between soft and stiff microgels suggesting that an increase in nascent matrix for (other) proteins is not inherent to softer microgels (Figure 3h). The chondrogenic master transcription factor SOX9 was increased nearly threefold in fluorescence intensity in soft microgels compared to stiff microgels (Figure 3i). Moreover, MSCs in soft microgels had a significantly higher nuclear/cytoplasmic ratio of SOX9 than in stiff microgels (Figure 3j). This suggests that the mechano-stimulated chondrogenesis of MSCs that are phenol-tethered within soft microgels occurred over the classical chondrogenic SOX9-mediated signaling axis. Together, this revealed that a pericellular environment as soft as 14 kPa Young’s modulus stimulated chondrogenesis, while microenvironments as stiff as 38 kPa did not augment chondrogenesis, thus confirming that covalent tethering of cells to materials via phenolic crosslinking mediates mechanotransduction to guide stem cell fate including chondrogenesis.

**Figure 3:**
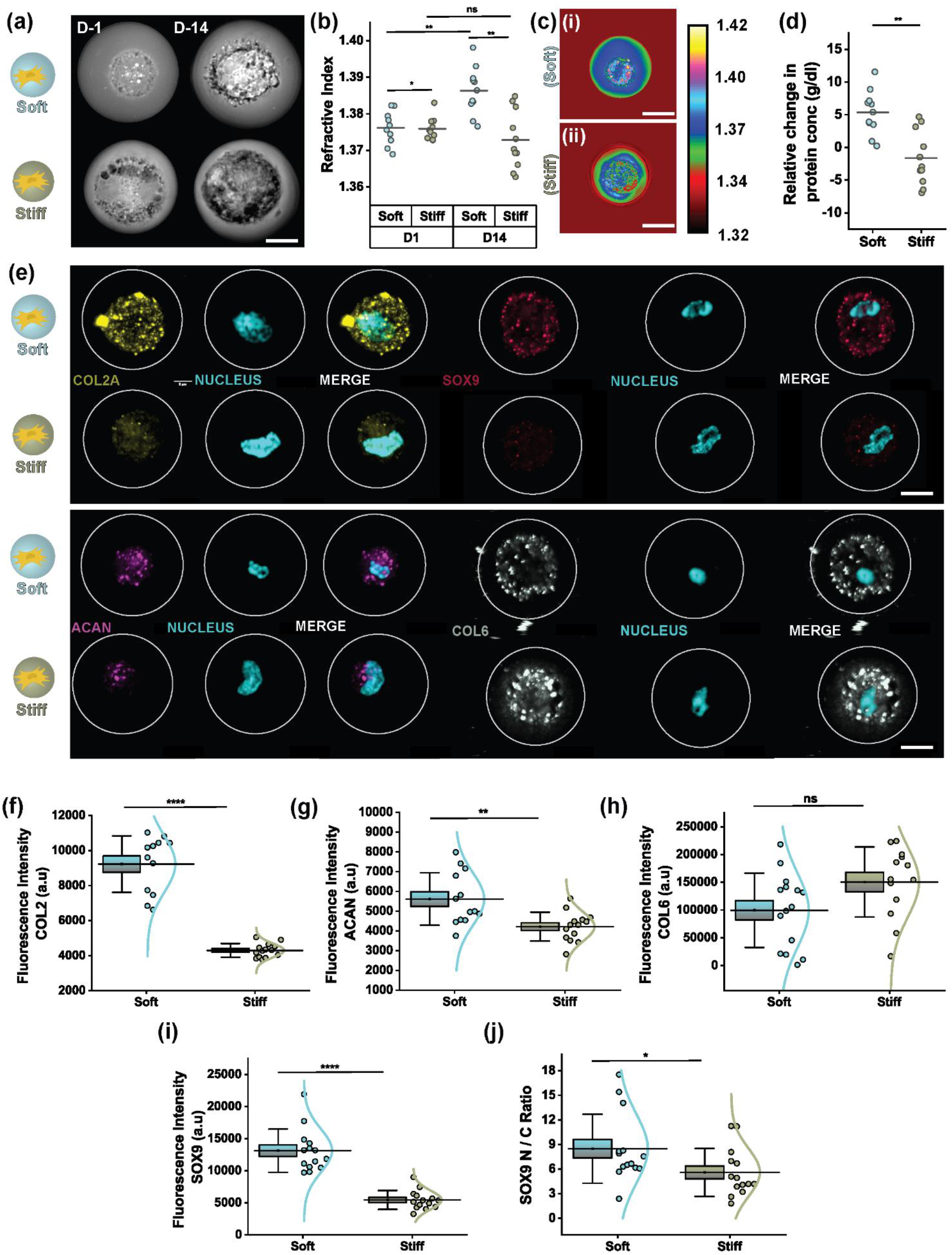
Mechanotransductive guidance of chondrogenic stem cell fate via oxidative phenolic crosslinking within single-cell microgels. **(a)** Holotomographic QPI imaging (Scale bar indicates 5 µm) and **(b)** refractive index quantitation of soft and stiff single-cell microgels after 1 and 14 days of chondrogenesis. n≥10. **p<0.01, *p<0.05 by Kruskal-Wallis ANOVA. **(c)** Holotomographic micrographs showing deposited protein intensity distribution in (i) soft microgels (ii) and stiff microgels (Scale bar indicates 5 µm). **(d)** Quantification for protein concentration after two weeks of chondrogenesis from soft and stiff microgels. n≥10, **p<0.01 by Kruskal-Wallis ANOVA. **(e)** Fluorescence confocal micrographs of soft and stiff single-cell microgels stained for chondrogenic differentiation markers COL2 (yellow), ACAN (magenta), COL6 (gray), and chondrogenic master transcription factor SOX9 (red) (Scale bar indicates 5 µm). Image-based quantitation of total fluorescence intensity in soft and stiff single-cell microgels of **(f)** COL2, **(g)** ACAN, **(h)** COL6, **(i)** SOX9, and **(j)** relative Nuclear/Cytoplasmic ratio of SOX9. n≥ 13, ****p<0.0001, **p<0.01, *p<0.05 by Kruskal-Wallis ANOVA. Box plots depict the standard error of the mean, and whiskers depict the standard deviation.

### Chondrogenic lineage commitment via oxidative phenolic crosslinking of MSCs in microgels is independent of YAP/TAZ-mediated mechanotransduction

Conventional stimulation of chondrogenesis via material stiffness (e.g., mediated via RGDs) operates over the YAP/TAZ signaling axis. To investigate whether mechanotransduction via covalent oxidative also involves classical YAP/TAZ signaling, we introduced a sequence of cyclic RGDs within Dex-TA microgels and compared these microgels to pristine Dex-TA microgels in terms of YAP/TAZ expression and chondrogenic differentiation. A 2D control of MSCs on a stiff surface was used to demonstrate the ability to detect YAP/TAZ expression and spatial localization (Figure 4b). While clear YAP nuclear translocation was observed in 2D, nuclear translocation was neglectable in 3D regardless of stiffness (Figure 4c) or the presence of RGDs (Figure S3). This suggested that within Dex-TA microgels - that facilitate mechanotransduction via oxidative phenolic crosslinking - YAP/TAZ signaling did not play a pivotal role, even when presenting RGDs. In sharp contrast, chondrogenesis of MSC in Dex-TA microgels was potently affected by the presence or absence of RGDs (Figure 4d). Specifically, while both COL2 and ACAN were significantly more expressed in soft microgels than stiff microgels regardless of RGD presence, the addition of RGDs significantly decreased the expression of COL2 (Figure 4e) as well as ACAN (Figure 4f). This finding validated that covalent oxidative phenolic crosslinking-mediated mechanotransduction facilitates chondrogenic lineage commitment of mesenchymal stem cells more efficiently than the conventionally explored RGD formulations. The addition of RGDs into the oxidative phenolic crosslinking formulation effectively inhibited the chondrogenic differentiation process, which supports the potency of oxidative phenolic tethering of cells to materials in terms of chondrogenic differentiation.

**Figure 4:**
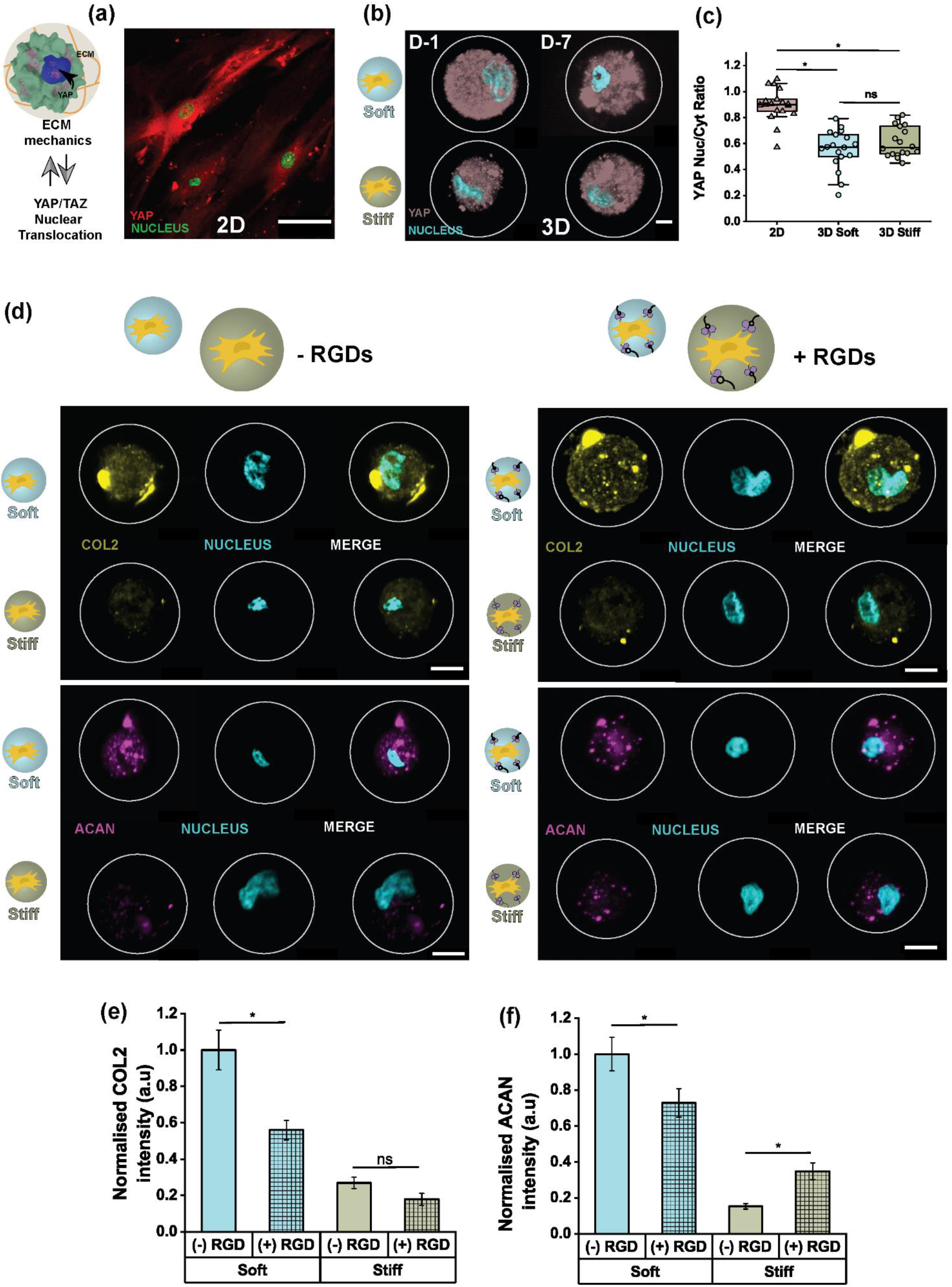
Adhesion RGDs and YAP/TAZ independent mechanotransduction of oxidative phenolic crosslinked single cell microgels. YAP/TAZ activity measurement of MSCs in microgels based on the matrix mechanics in differentiation state post one week after encapsulation. **(a)** Fluorescence confocal micrographs of YAP/TAZ (red) nucleus (green) stained MSCs that were adhered on TCP(Scale bar indicates 100 µm). **(b)** Maximum intensity projected 3D images of MSCs in microgels expressing YAP/TAZ (brown) nucleus (cyan) (n≥11 per condition) (Scale bar indicates 5 µm), **(c)** YAP/TAZ Nuc/Cyto ratio of MSCs on 2D and in microgels (soft and stiff) (*p<0.05 by Kruskal-Wallis ANOVA). **(d)** Comparison of fluorescence confocal images of soft and stiff microgels in the presence and absence of RGDs analyzed for expression of chondrogenic differentiation markers COL2 (yellow) ACAN (magenta) post two weeks of differentiation (Scale bar indicates 5 µm). Normalized fluorescence intensity of **(e)** COL2, **(f)** ACAN compared between microgels with and without RGDs (n≥20 per condition) (***p<0.001, *p<0.05 by Kruskal-Wallis ANOVA). (The box plots show 25/75th percentiles and the whiskers show the maximum and minimum data points. All data shown are mean± s.e.m

### Microscale mechanics induce mechanistic alterations in the cytoplasm during fate transitions

Sensed matrix mechanics are most commonly transduced via the cytoplasm by regulating various intracellular signaling events, which can influence stem cell behavior including differentiation^34^. Of specific interest, cytoplasm-associated mechanical adaptation can alter cell size, cytoplasmic volume, and nuclear volume, which are known to be able to influence stem cell function via largely unknown mechanisms^35^. To investigate whether oxidative phenolic crosslinking-mediated mechanotransduction could also potentially affect cell function by altering cytoplasmic volume, MSCs were encapsulated in either soft or stiff Dex-TA microgels, differentiated into the chondrogenic lineage, and quantitatively analyzed on cytoplasmic volume using real-time live cell confocal imaging (Figure 5a). This demonstrated that MSCs in both soft and stiff microgels underwent significant volume reductions over time (Figure 5b). Cell volume quantitation based on 3D reconstructed live cell fluorescence images corroborated this observation (Figure 5c). However, the extent of volume reduction at day seven was slightly different among cells in soft and stiff microgels. MSCs in stiff microgels showed a higher volume reduction (38%) as compared to their soft counterparts (30%) (Figure S4). To further validate if the volume reduction is due to differentiation, we investigated the cytoplasmic volume of MSCs in soft and stiff microgels when exposed to a proliferation medium. We observed there were no significant changes in volume for MSCs encapsulated in either soft or stiff microgels over time (Figure S4). Hence, the observed volume reductions were specific to differentiating single cells irrespective of their stiffness as previously reported^27^. Even though the volume reductions were not highly stiffness dependent, interestingly, alterations in cell volume can cause altered intracellular molecular crowding and thereby influence cellular function^36^, which has not yet been explored in 3D microenvironments, especially not in the context of the effect of phenolic crosslinking-induced cell volume reduction. We hypothesized that distinct microgel stiffness together with cell volume could influence intracellular molecular crowding. To validate this hypothesis, we investigated the dry mass and cell volume that were highly correlated to each other in both soft and stiff microgels (Figure S5).

**Figure 5:**
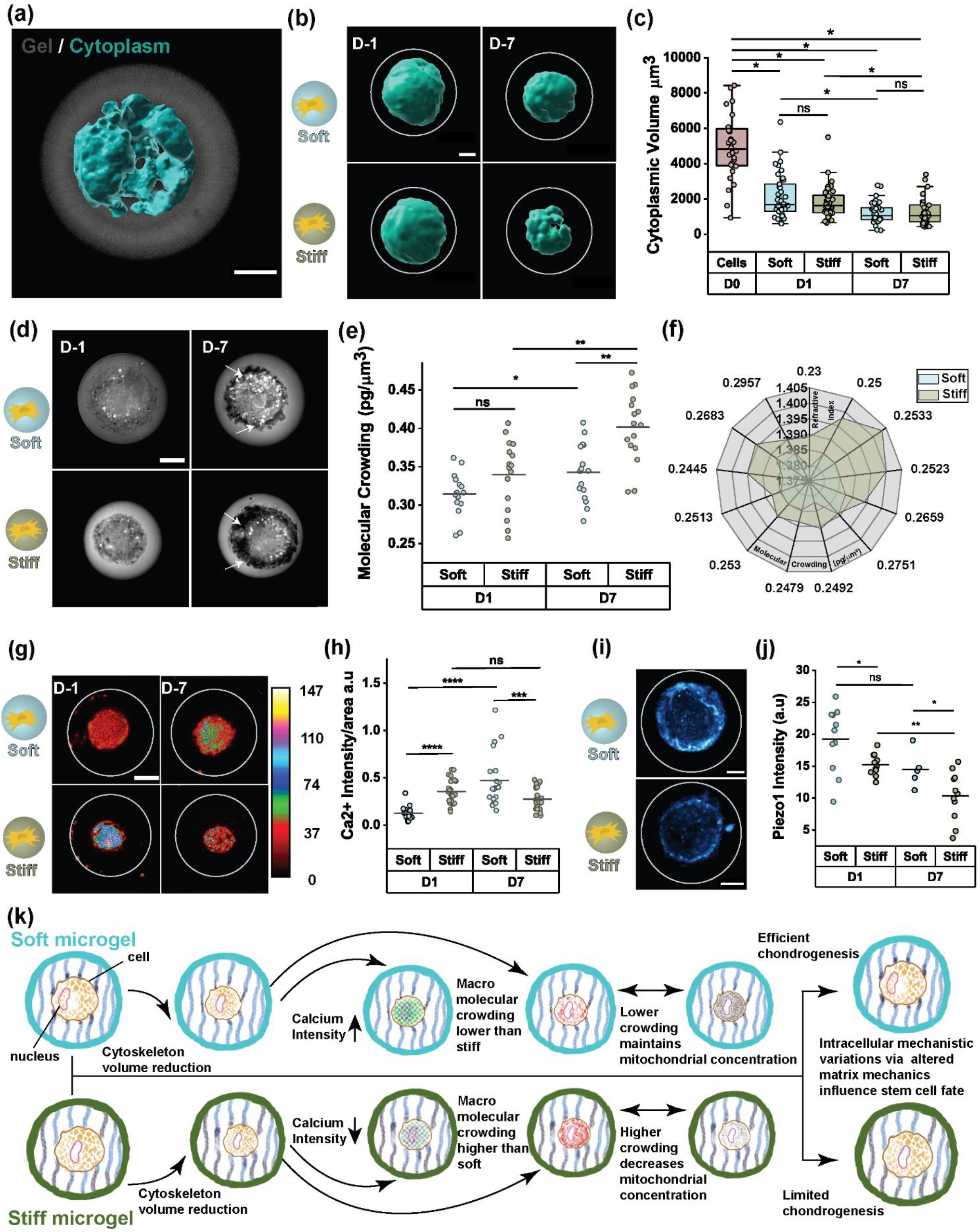
Stiffness-dependent intracellular mechanistic changes in stem cells via oxidative phenolic mechanotransduction. **(a)** 3D reconstructed cytoplasm of a single cell inside a microgel (Scale bar indicates 5 µm). **(b)** 3D reconstructed temporally mapped confocal images using IMARIS from soft and stiff microgels also denote volume reduction was stiffness dependent (n≥31) (Scale bar indicates 5 µm). **(c)** Cytoplasmic volume quantification of MSCs in soft and stiff microgels during differentiation for 7 days (n≥31) (*p<0.05 by Kruskal-Wallis ANOVA) The box plots show 25/75th percentiles and whiskers show the maximum and minimum data point. **(d)** Holotomographic QPI imaging of MSCs in soft and stiff microgels to estimate the molecular crowding up to 7 days of differentiation (Scale bar indicates 5 µm). **(e)** Quantification of molecular crowding calculated from dry mass and volume per single MSC in soft and stiff microgels up to 7 days during differentiation (n≥15) (**p<0.01, *p<0.05 by Kruskal-Wallis ANOVA). **(f)** Refractive index mapping as a measure of molecular crowding in soft and stiff microgels after 7 days (n≥15). **(g)** Representative confocal images of intracellular calcium intensity in cells on soft and stiff microgels. **(h)** Quantification of intracellular calcium intensity over measured area of MSCs in soft and stiff microgels at D1 & D7 (n≥19 per condition) (**** p<0.0001, ***p<0.001, **p<0.01 by One-way ANOVA) (Scale bar indicates 5 µm). **(i)** Fluorescence representative images and **(j)** quantification, of cells stained for PIEZO1 in soft and stiff Microgels after 7 days (n≥5). (*** p<0.001, ns – not significant) by Kruskal-Wallis ANOVA) (Scale bar indicates 5 µm). **(k)** Schematic representation of summarizing events that occurred in soft and stiff microgels depicting stiffness-driven volume reduction influences intracellular molecular crowding and calcium signaling affecting mitochondrial activity thereby potentially influencing stem cell fate of MSCs inside soft and stiff microgels.

Consequently, it was revealed that the molecular crowding in MSCs in stiff microgels was significantly higher than that of MSCs in soft microgels, and these differences became larger over time (Figure 5d, e). To further investigate this observation, we analyzed the refractive index changes as a measure of increased molecular crowding. A higher refractive index value corresponds to a higher molecular crowding as reported previously^32^. Indeed, MSCs in stiff microgels showed a higher refractive index value corresponding to a higher intracellular crowding than MSCs in soft microgels (Figure 5f). It has previously been reported that volumetric compression can elevate intra-cellular crowding in intestinal stem cells and maintain higher proliferation thereby hindering differentiation^37^, which complements our findings where MSCs in stiff microgels undergo higher volumetric compression and elevated intracellular crowding thereby having hindered differentiation potential. Such volume changes and altered molecular arrangements in the cytoplasm are largely influenced by water efflux, which potentially was anticipated to alter the calcium ion concentration, which balances the osmolarity ^38^. In this context, we investigated the distribution and concentration of calcium ions in MSCs in soft and stiff microgels (Figure 5g). In previous studies on cells on 2D surfaces that adhere based on dynamic reversible bonding moieties, stiffness increased intracellular calcium intensity in a stiffness-dependent manner^39^. However, how this relationship operates in 3D has remained poorly understood, we here reveal that the calcium intensity in MSCs in soft microgels strongly increased over time, while a mild yet significant reduction in calcium intensity occurred in stiff microgels (Figure 5h). We hypothesized, that fluctuations in calcium ion concentration might be achieved via mechanosensitive ion channels. Hence, we next investigated the activity of Piezo1, a mechanosensitive ion channel that mediates stretch-activated currents in chondrocytes^40^ and inhibits stem cell fate^41^. We observed that Piezo 1 was significantly higher in MSCs inside soft microgels compared to stiff microgels, which was consistent over time (Figure 5i & 5j). Moreover, Piezo1 is known to regulate mitochondria^42,43^, and we indeed also noted a reduction in the mitochondrial concentration within MSCs in stiff microgels as compared to soft microgels over time (Figure S6). Collectively, these findings provide a interlinked and multifaceted yet streamlined mechanism via which covalent bonding of cells to their microenvironment via oxidative phenolic crosslinking can achieve mechanotransduction and subsequent orchestration of stem cell function based on stiffness dependent cytoplasmic volume modulation (Figure 5k).

### Oxidative phenolic crosslinking mediates mechanotransduction by altering nuclear organization

In addition to cytoplasmic changes, nuclear volume changes can also cause changes in cellular behavior. For example, it has been reported that biophysical alterations such as volume and aspect ratio changes to the nucleus can alter protein production, gene transcription, and chromatin localization^44^. However, whether this mechanism could play a role in stiffness-induced lineage commitment of stem cells with 3D microenvironments or via oxidative phenolic crosslinking has remained unknown. Hence, we investigated whether culturing MSCs in either soft or stiff Dex-TA microgels altered their nuclear volumes. To this end, we leveraged live single-cell nuclear imaging to effectively discriminate between microgel, cytoplasm, and nucleus in a voxelated manner, which allowed for the quantitative determination of single nucleus volume (Figure 6a). 3D reconstruction on MSCs in soft and stiff Dex-TA microgels revealed that while nuclear shape (e.g., ellipticity) remained similar (Figure S8), the nuclear volume of MSCs was altered in a stiffness-dependent and time-dependent manner (Figure 6b). Specifically, nuclei of MSCs in stiff microgels had shrunk and then increased significantly as compared to those cultured stiff microgels one day post-encapsulation. (Figure 6c) and also these changes were specific to differentiation (Figure S4). We correlated whether cytoplasm-induced volume changes had effects on nuclear volume, surprisingly, the cytoplasm and nuclear in soft microgels had a negative correlation while stiff microgels showed a positive correlation (Figure S7). This validated that pericellular mechanics sensed via oxidative phenolic crosslinking also was transduced to the nucleus. Interestingly, the cytoplasm and nucleus operate biophysically distinct and respond to the sensed mechanics in highly a stiffness-dependent manner.

**Figure 6:**
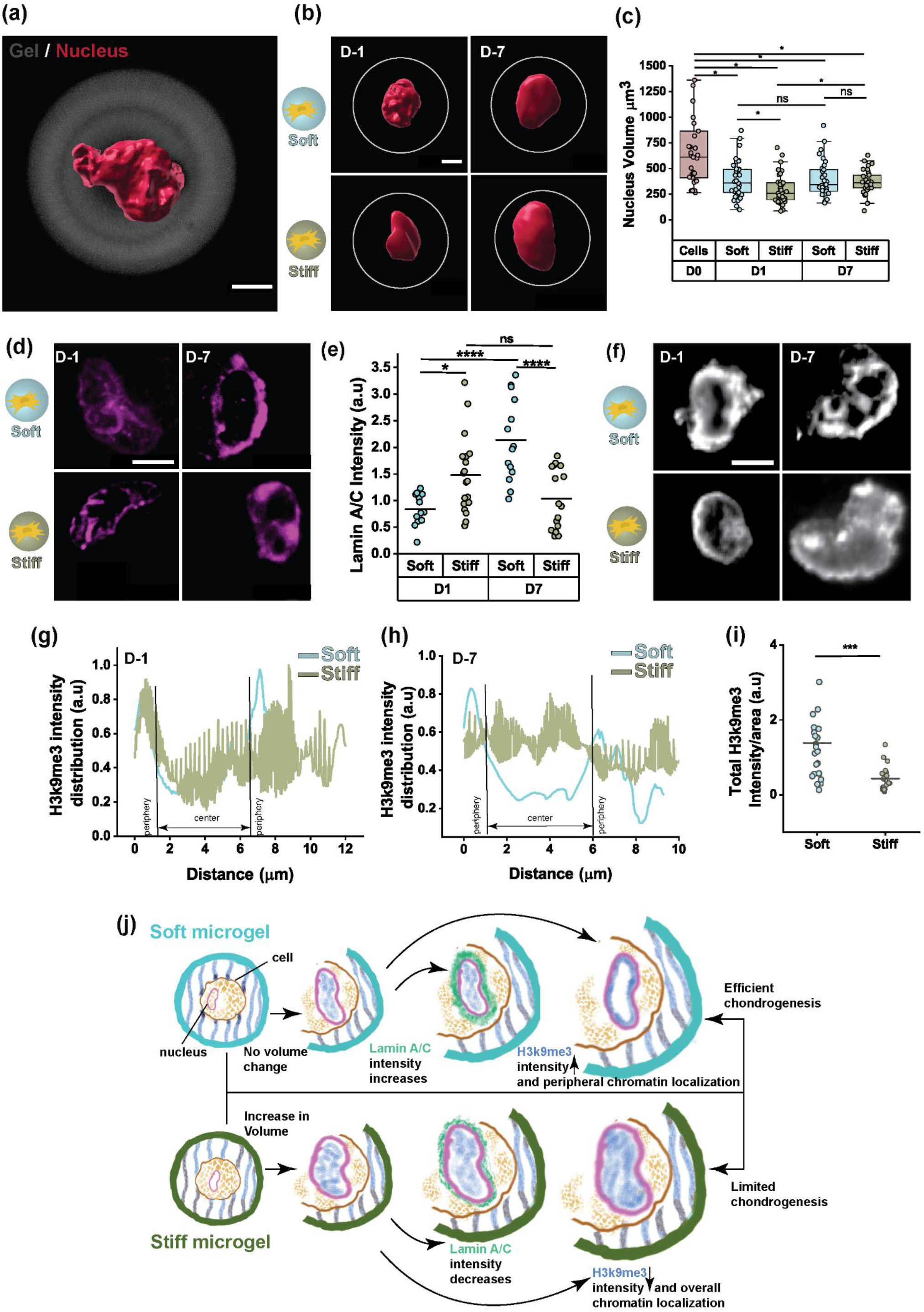
Stiffness mediated nuclear mechanoresponse of single MSCs in oxidative phenolic crosslinked microgels. **(a)** 3D reconstructed nucleus of a single-cell inside a microgel (Scale bar indicates 5 µm). **(b**) 3D reconstructed representative confocal images at day 1 (D1) & day 7 (D7) of single MSCs’ nuclei in soft or stiff microgels (n≥31 per condition) (Scale bar indicates 5 µm). **(c)** Nucleus volume quantification of cells in soft and stiff microgels during differentiation up to 7 days (n≥31 per condition) (*p<0.05 by Kruskal-Wallis ANOVA). Box plots show 25/75th percentiles and whiskers show the maximum and minimum data points. **(d)** Fluorescence confocal micrographs (Scale bar indicates 5 µm). and **(e)** image-based intensity quantification of nuclear Lamin A/C expression of MSCs from the soft and stiff microgels up to 7 days(n≥13) (**** p<0.0001, *p<0.5 by one-way ANOVA). **(f)** Fluorescence micrographs of MSC nuclei stained for H2k9me3 in soft and stiff microgels for up to 7 days (n≥22 per condition) (Scale bar indicates 5 µm). Image-based H3k9me3 intensity distribution in soft and stiff microgels during **(g)** day 1 (D1)and **(h)** day 7 (D7) of differentiation (n≥10). **(i)** Total H3k9me3 intensity of single nuclei of MSCs in both soft and stiff microgels after 7 days of induced differentiation (n≥22 per condition) (***p<0.001 by Kruskal-Wallis ANOVA). **(j)** Schematic drawing depicting the mechanoresponse of the nucleus to altered pericellular matrix mechanics affecting nuclear volume, Lamin A/C protein production, and H3k9me localization contributing to transitions in stem cells.

To investigate whether this stiffness-induced nuclear alteration was mediated via a direct (e.g., nuclear mechanical forces) mechanism, we studied the expression of nuclear Lamin a/c, which informs on the stress on the nuclear envelope. As changes in Lamin a/c require time to visualize and stabilize, we stained MSCs in soft and stiff microgels for up to seven days of culture during chondrogenic differentiation (Figure 6d). Lamin a/c expression showed a trend of being higher in stiff microgels compared to soft microgels one-day post-encapsulation, which aligned with the more intense reduction in nuclear volume in stiff microgels. Interestingly, after seven days in differentiation culture, MSCs in soft microgels had significantly upregulated their Lamin a/c expression, while those of MSCs in stiff microgels had remained relatively stable (Figure 6e). To determine whether these observations could be involved in altered cellular function, we next investigated the chromatin organization of the nuclei, which can be regulated by altered nuclear stress^45,46^. MSCs in oxidative phenolic crosslinked soft or stiff Dex-TA microgels were investigated on their expression patterns of H3K9me3 (Figure 6f), which is a canonical histone tail modifier known to alter cellular function via transcriptional silencing. Total H3k9me3 stain intensity showed an intimate correlation with Lamin a/c expression, as H3k9me3 was higher in MSCs in soft microgels than stiff microgels at seven days post-encapsulation (Figure 6i). These results complemented previously reported findings where Lamin a/c expression corroborated with H3k9me3 methylation in MSCs in varying nano topographies^47^. In addition, analysis of H3k9me3 spatial distribution patterns revealed that MSCs in soft microgels were characterized by a high level of chromatin at their nuclear periphery, while the chromatin distribution of MSCs in stiff microgels was less intense and more randomly distributed inside the nucleus (Figure 6g & h). These results align with previously reported studies where substrate stiffness regulates chromatin condensation and localization in hMSCs and interferes with stem cell differentiation^48^ Collectively these findings imply that the nucleus has a distinct mechanoresponse to matrix mechanics being transduced via oxidative phenolic crosslinking, which displays a stiffness dependent chromatin organization (Figure 6h). Together, this demonstrates that mechanotransduction via oxidative phenolic crosslinking can directly convey nuclear stresses, which can alter cellular behaviors and potentially alter or drive cellular fate.

## Conclusion

Herein, we explore and explain the mechanotransduction mechanism of covalent on-cell material tethering, and that of oxidative phenolic crosslinking mediated lineage commitment of stem cells in particular. This study yields key mechanistic insights and an understanding of how the covalent bonding of biomaterials to single cells orchestrates and influences stem cell differentiation in a stiffness-dependent manner. Interestingly, we demonstrated that the introduction of commonly used adhesive ligands such as RGDs can adversely affect the mechanotransduction-mediated chondrogenic commitment of MSC that are tethered to biomaterials via on-cell oxidative phenolic crosslinking, which suggests that direct covalent tethering of materials to cell membranes can offer superior control over lineage commitment, at least for chondrogenesis. In contrast to conventional non-covalent dynamic cell binding strategies, mechanotransduction in covalent on-cell reacted materials was independent of conventional YAP/TAZ pathway and instead was mediated via biophysical alterations of intracellular organization. By leveraging a microfluidically driven single-cell encapsulation system, we were able to track the mechanistic changes in the MSCs’ intracellular behavior at single-cell resolution in a high throughput order. Intriguingly, the cytoplasmic changes led to various cascades of molecular changes correlating with nuclear changes that were linked to chromatin organization and altered protein production, all dependent solely on the stiffness of the pericellular matrix. Hence, we report that covalent tethering of cells to materials offers a novel opportunity for mechanistic control over cellular function and fate, which can be used for fundamental biological studies, cellular therapeutics, drug testing, lab-grown meat, fabrication, and tissue engineering applications.

### Experimental Section

#### Polymer Synthesis

Dextran (40kDa, Pharmacosmos, Denmark) was functionalized with tyramine (Sigma Aldrich) as previously reported. In short Dextran (5g) with Lithium chloride (Sigma Aldrich) 4g was transferred to a round bottom flask which is maintained at 90°C, and moisture is removed by repeated flushing of nitrogen and air up to 3 cycles. After which 200ml of Dimethylformamide (Sigma Aldrich) (DMF, 99.8% anhydrous) is added to the flask under an inert atmosphere and constant heating up to 90°C. When the solution turned transparent the mixture was cool down to 0°C and previously purified 4-nitrophenyl chloroformate (PNC, 96%) (Sigma Aldrich) was added in steps maintaining the inner temperature below 2°C. The reaction was carried out for 1 hour and the resulting product was precipitated in cold ethanol and diethyl ether (VWR), vacuum-filtered and oven-dried. Subsequently, the dried product was dissolved in 200ml of DMF followed by the addition of tyramine and was left for 1 hour. The resulting solution was precipitated in cold ethanol and diethyl ether, vacuum filtered, and dialyzed for 4 days using a (Spectra/Por) 1kDa MWCO dialysis membrane. The freeze-dried obtained white powder was analyzed using 1H-NMR which confirmed successful functionalization with 15 tyramine moieties per 100 repetitive dextran monomer units.

#### Tethering experiment

To first demonstrate tethering tyramine-tyrosine crosslinking was established and analyzed as previously reported. To verify, we analyzed samples using a UV-vis spectrophotometer (Agilent Technologies Cary 300 UV-vis) to determine relative peak shifts. Briefly, 0.5mM tyramine and tyrosine (Sigma Aldrich) solutions in PBS were mixed with 3U/ml HRP and 0.03 w/v% of H_2_O_2_ (Sigma Aldrich) and reacted for 5 minutes before transferring into a quartz cuvette for measurement, H_2_O was used as a substitute for H_2_O_2_ in negative control samples. To confirm oxidative phenolic tethering on cell membranes, 0.2ul of 100x AF647-Tyramide (Alexa Fluor 647, Thermo Fisher) solution was added to cells (10.000 cells/ml) suspended in a cell culture medium. After incubating for 15 minutes, cells were washed twice to remove excess AF647 reagent and visualized using confocal microscopy (Zeiss LSM 880). To demonstrate tethering at the material interface, 10w/v% DexTA, 10U/ml HRP, and 0.03% H_2_O_2_ were used to produce hydrogel discs of 3mm diameter using PDMS molds. The washed crosslinked hydrogel was then incubated with AF647-tyramide reagent, 3U/ml HRP, and 0.03% of H_2_O_2_, crosslinked for 3 minutes, and washed twice with PBS before being imaged using a confocal microscope. To visualize cell-material tethering, cells labeled with Mitotracker green (Fischer M7514) were seeded on top of hydrogel discs with AF647, prepared as mentioned above. Tethering was achieved by adding 44U/ml of HRP and 0.03% of H_2_O_2_ to react for 90 seconds and washed immediately with PBS. Discs were imaged using brightfield and confocal microscopes to visualize the tethering of cells on hydrogel discs.

#### Fabrication of microfluidic platforms

The microfluidic devices were manufactured using standard soft lithography techniques. Briefly SU8-3025 (Kayaku) was poured onto one-side polished silicon wafer and cured using UV light exposed through a patterned mask with distinct structures. The obtained silicon wafers were filled with polydimethylsiloxane (PDMS) mixed in a 10:1 ratio with the crosslinkers, degassed, and cured at 60 °C overnight. The negative of the structures obtained on the PDMS was bonded to the glass slide using an oxygen plasma cleaner (FEMTO Science Inc, South Korea) and kept at 60 °C overnight for attachment of PDMS to glass. The produced microfluidic chips had an approximate height of 25µm (droplet generator chip) and 100µm (crosslinking chip). All microfluidic chips were Aquapel (Amazon) treated in the channels before usage to ensure channel wall hydrophobicity.

#### Cell culture

Mesenchymal stem cells (MSCs) were isolated from bone marrow tissues obtained from patients with written consent and approved by the ethical committee of the Medisch Spectrum Twente. (METC\06003). Cells isolated from the human tissues were expanded at a density of 2800 cells/cm^2^ cultured in an MSC proliferation medium consisting of 10v/v % FBS, 100mg/ml penicillin, 100mg/l streptomycin (Gibco), 0.2 x 10-3 M ascorbic acid (Sigma Aldrich), 1v/v% Glutamax (Gibco), 1ug/l bFGF (Neuromics) (added every time fresh) in aMEM (GIBCO) at 37°C under 5% CO_2_. The medium was replenished two times a week before attaining near confluency after which harvested using 0.25% v/v Trypsin-EDTA at 37°C. The harvested cells were expanded (up to passage 5 maximum) further or used to produce cell-laden microgels.

#### Production of microgels

Aquapel treated chips were mounted on a custom-built PMMA platform and connected to gastight syringes (Hamilton) using FEP tubing, which was controlled by low-pressure (Cetoni) syringe pumps. The droplet generator had a continuous phase containing Novec 7500 (Flurochem) engineered oil with 2% Picosurf (Spherofludics) as a surfactant. The dispersed phase consisted of DexTA 10 w/v%, HRP (Sigma Aldrich 25kU) 44U/ml Optiprep 8% (Sigma Aldrich), and mesenchymal stem cells (MSCs) 10^7^ cells/ml. In the dispersed phase when flow focused into the continuous phase, emulsions were formed using flowrates 6:2 (µl/min) (oil: polymer), which then was in-line flown into a crosslinking chip through which H_2_O_2_ was flowing through parallel channels in the opposite direction as the microemulsion. The crosslinker feed concentration used was 2.5w/v% (soft) and 10w/v% (stiff). The diffusion-based crosslinking enabled the formation of crosslinked microgels around single cells. The collected microgels were washed and retrieved from the oil phase using Picobreak (Spherofluidics) and then induced into chondrogenic differentiation medium consisting of 100mg/ml penicillin, 100mg/l streptomycin (Gibco15140148), 1x ITS-premix (Corning), 40ug/ml proline, 50ug/ml ascorbic acid, 1x sodium pyruvate (Sigma Aldrich), 10ng/ml TGFbeta3 (R&D systems) and 10^-7^ M dexamethasone (Sigma Aldrich) added fresh in DMEM (Gibco). The medium is refreshed thrice a week for up to 14 days of differentiation.

#### Nanoindentation of microgels

Microgels were characterized for their mechanical property using Pavone (Optics 11 life) nanoindenters. Briefly, the microgels were added to non-treated cell culture plates, which were treated with 2mg/ml dopamine (Sigma Aldrich) in TRIS buffer (Sigma Aldrich) for 60 minutes, and washed thoroughly before the addition of microgels to prevent mobilization during measurements. All measurements were carried out in PBS unless when microgels with cells were measured, which was carried out in a differentiation medium. The cantilever with a spring constant (0.25 N/m) consisted of a spherical tip of radius 11um which was brought in contact with the surface of the microgel. The indentation control mode was selected with a sample indentation depth of 400nm for every microgel indenting for 2s followed by a hold period of 1s and 2s for retraction. The obtained indentation curves were fitted using a Hertzian model from the Data viewer software (Optics 11 Life). The elastic modulus (referred to as Youngs Modulus) was calculated by E= Eeff (1-𝜈2) where Eeff is the effective Youngs modulus, E is the Youngs modulus and 𝜈 is the Poisson ratio (assumed as 0.5 for hydrogels). For DMA measurements, the above-mentioned settings were maintained and indentation was carried out at frequencies varying from 1Hz to 10 Hz.

#### Live Dead and Mitochondrial activity

Live/Dead analysis of cells was performed by incubating single cell microgels in 2 x 10^-6^ calcein (live) and 2 drops/ml of NucBlue (nucleus) (Thermo Fisher) reagent for 30 minutes and visualized using fluorescence microscopy. Cell viability was determined as a fraction of live cells (presence of calcein signal) divided by the total number of cells. To determine the mitochondrial activity 5um of MitoTracker Green FM (Thermo Fisher) solution was incubated along with the Microgels for 60minutes washed three times with PBS before fluorescence imaging using EVOS (Thermo Fisher)

#### Confocal Fluorescent Imaging

To visualize the crosslinking density, acellular microgels were incubated with 4 x 10^-6^ M EthD-1 for 30 minutes and visualized under 10x objective with N/A confocal microscope (Zeiss LSM 880 airy scan). For diffusion studies, the microgels were incubated in FITC-Dextran 20kDa (Sigma Aldrich) at 1mg/ml in PBS and time-resolved confocal imaging was performed for 24 hours to monitor the movement of FITC-Dextran into the microgels.

For immunofluorescence imaging to determine chondrogenic differentiation markers, single-cell microgels were cultured in differentiation medium for two weeks, fixated in 4% formaldehyde solution (Sigma Aldrich) (Sigma Aldrich) for 15 minutes washed twice with PBS, and stored at 4°C before usage. Before imaging, single-cell microgels were permeabilized using 0.25 w/w% Triton-X 100 (Sigma Aldrich) in PBS for 30 minutes washed two times with PBS, followed by blocking using 1% BSA (bovine serum albumin) (Sigma Aldrich) for 60 minutes. Microgels were washed twice in 1x DPBS before incubating with primary antibody for anti-collagen II (Abcam ab34172, 1:100) anti-aggrecan (Abcam ab3778, 1:100), anti-collagen VI (Abcam ab182744, 1:100), anti-SOX9 (Novus Biologics NBP1-85551, 1:100) anti-YAP (Santa Cruz sc101199, 1:50), anti-Lamin A/C (Santa Cruz sc7292, 1:25), anti-PIEZO1 (Novus Biologics NBP1-78446, 1:100), anti-H3k9me3 (Abcam ab8898, 1:50). All dilutions were performed in BSA and incubated at 4°C overnight. Before the addition of the secondary antibody, microgels were washed three times with 1x PBS and incubated with goat anti-rabbit Alexa Fluor 488 and goat anti-mouse 647 accordingly overnight at 4°C, and washed three times with 1x DPBS. The nucleus was labeled with DAPI (Thermo Scientific 62248 1:100), incubated at RT for 15 minutes, and washed with 1x DPBS before suspending the microgels in Ibidi µslide uncoated (81821) for imaging. The fluorescence imaging was performed using a 63x w/o objective NA 1.2 (Zeiss LSM 880 airyscan) microscope and obtained images was processed with ImageJ and IMARIS software.

For calcium imaging, cells in microgels were incubated in 5uM of fluo-4AM for 60minutes, and the nucleus was labeled with 2 drops/ml of NucBlue (Thermo Fisher), and incubated for 15 minutes, washed three times with 1xDPBS before imaging. The intracellular calcium intensity was imaged every second up to 20s for each microgel individually. The thickness of the microgel area was measured using actin cytoskeleton staining to determine the cell membrane boundary and the microgel was stained using EthD-1. Briefly, microgels were fixated in 4% formaldehyde solution for 15 minutes and washed twice with PBS before permeabilization using Triton-X 100 for 30 minutes. Blocking with 1% BSA for 60 minutes followed by incubation with Alexa Fluor 647 Phalloidin (1:40) was carried out, washed three times with 1x DPBS followed by nucleus labeling with DAPI (1:100) for 15 minutes before imaging using confocal microscopy. For cell volume analysis, cells in microgels were incubated in 2 x 10^-6^ calcein (live) and 2 drops/ml of NucBlue (nucleus) (Thermo Fisher) reagent for 30 minutes, and z-stack images were acquired on a (Zeiss LSM 880 airyscan) confocal microscope using 63x w/o objective NA 1.2 using two different channels.

#### Image Analysis

The immunofluorescence images for chondrogenic differentiation markers were analyzed using ImageJ and auto-thresholding was applied before measuring the mean intensity value for specific markers. For cytoskeletal and nuclear volume measurements, acquired z-stack images were 3D reconstructed using IMARIS by constructing surfaces on respective channels to obtain a 3D reconstructed model, volume and ellipticity were obtained as software-provided values. 3D YAP/TAZ nuclear to cytosolic ratio was calculated based on the following formula:

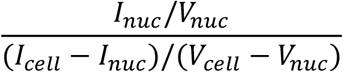

Where I_nuc_ and I_cell_ measured the total intensities of the YAP/TAZ signal inside the nucleus and the cell, and V_nuc_ and V_cell_ are the volumes of the nucleus and the cell measured from 3D reconstructed images from IMARIS. The calcium intensity of the cells in microgels was calculated based on a modified method as reported previously

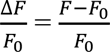

where F_0_ and F are the mean fluorescence intensities at T_0_ and T_20_ seconds The obtained value for divided over the measured area to determine the calcium intensity per cell per unit area.

#### Holotomographic QP imaging

Holotomography QP imaging was performed on HT-X1 (Tomocube Inc) to determine the protein deposition inside microgels. Briefly, after being chondrogenically differentiated for two weeks, microgels were fixed, suspended in PBS, and transferred to Tomodish provided by the manufacturer (a thin coverslip-enabled petri dish). The obtained images were processed using equipment-assisted TomoAnalysis software. RI values were used to determine the protein concentration in microgels using an adapted version of a previously reported method where *n(cell) = nm* + *αC(cell),* where *n(cell)* is the mean RI distribution of the cell, *nm* is the RI value of the surrounding medium (DPBS) (*nm* = 1.337at *λ* = 532 nm), *α* is an RI increment (*α* = 0.190 mL/g for protein), and C(cell) is the protein concentration of the cell. The data reported were from MIP images obtained from 3D holotomography. The dry mass and volume of the cell were obtained from the TomoAnalysis software by thresholding the cell regions from the microgel region thereby obtaining dry mass and volume of cells inside microgels. The extracted values are used to calculate the molecular crowding inside cells.

#### Data Analysis

All data analysis and statistical analysis were done using OriginPro 2019 software. The image processing was performed using ImageJ and IMARIS (9.9.1). The collected data were processed using OriginPro 2019. The drawings and schematics were prepared using Adobe Illustrator and Procreate.

## Acknowledgments

The authors thank Dr. Minye Jin (University of Twente) for assistance with the UV spectrometer measurement and analysis. Tom Knop (Bioimaging Centre, University of Twente) for assistance using confocal and holotomography.

