## Supplementary Info for "Covalent on-cell conjugation of biomaterials through oxidative phenolic coupling regulates stem cell fate via intracellular biophysical programming": Covalent on-cell conjugation of biomaterials regulates stem cell fate via intracellular biophysical programming_Supporting Information.pdf

**(a)**

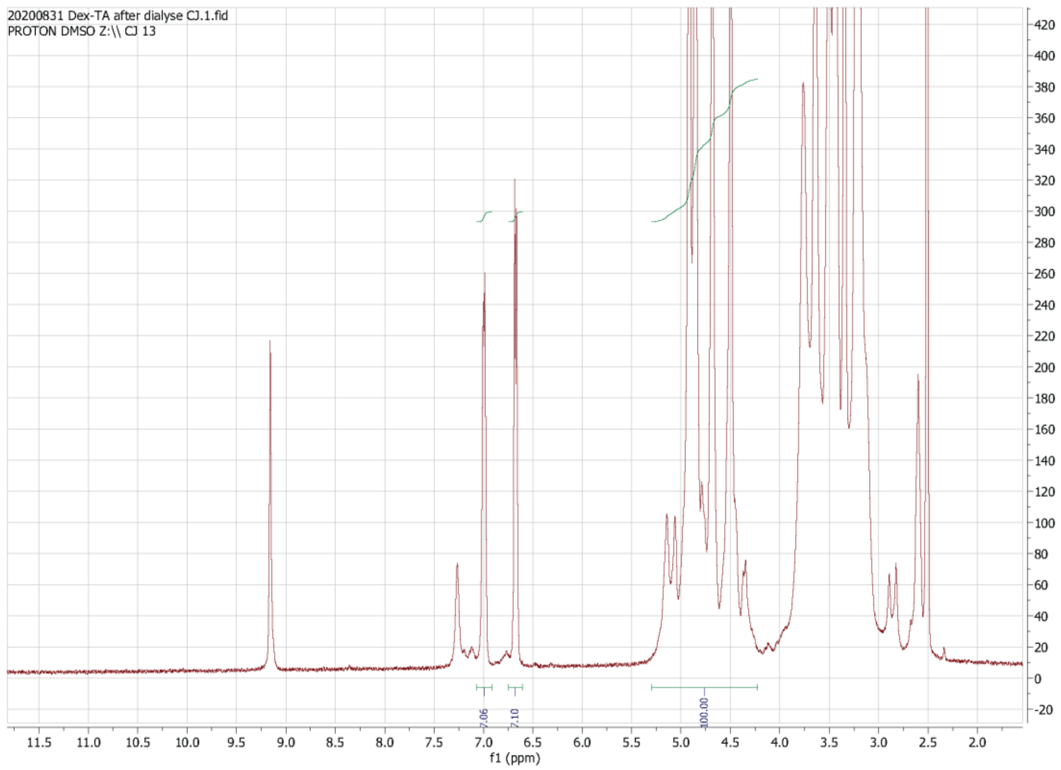


***Figure S1: (a)** [1]H-NMR of Dex-TA: The calculation of the DS of Dextran-PNC is based on integrals of fixed regions. The region of 4.2-5.8 ppm corresponds to 4 protons from dextran, while 7.4-7.65 and 8.2-8.45 together correspond to 4 protons from the (phenyl of the ) PNC. Setting the integral of 4.2-5.8 ppm to 100 directly results in the percentage of PNC by summating the integrals of 7.4-7.65 and 8.2-8.45.*

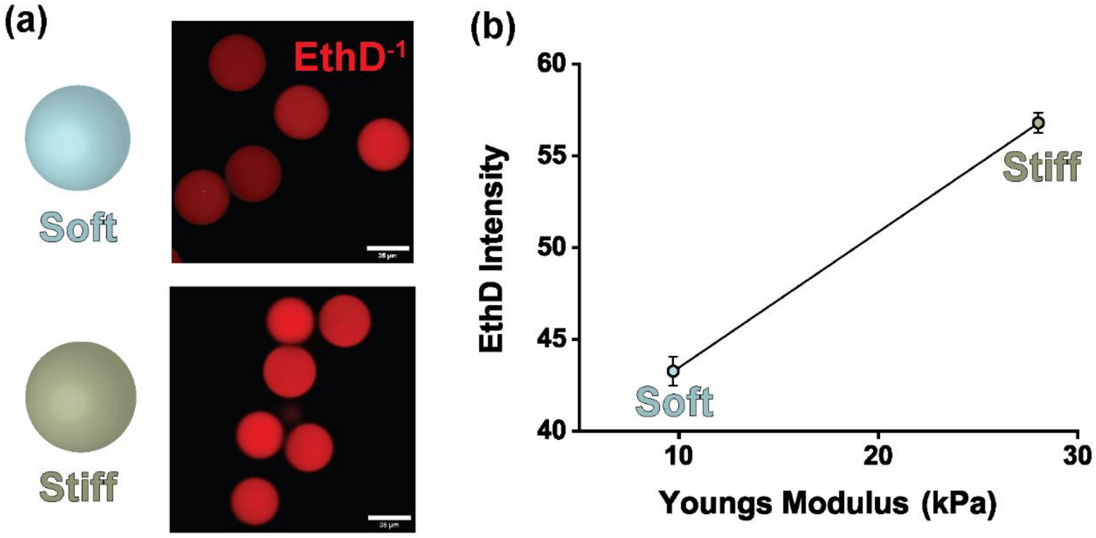


**Figure S2: (a)** *Fluorescence micrographs of ethidium homodimer (red) stained soft and stiff microgels* **(b)** *Fluorescence intensity of microgels correlated to Youngs Modulus of the microgels. (n≥35). Scale bar indicates 35µm.*

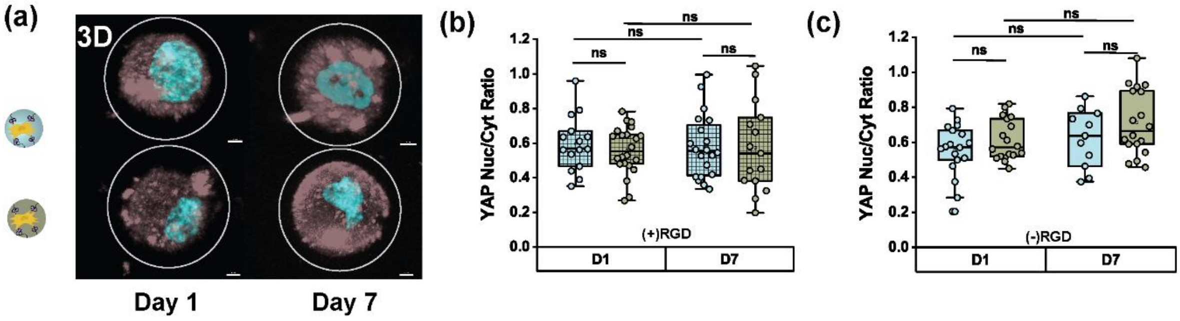


**Figure S3: (a)** *Maximum intensity projected 3D micrographs of microgels expressing YAP/TAZ (brown) nucleus (cyan) with RGDs. YAP/TAZ Nucleus-Cytoplasm ratio quantification from 3D confocal images from soft and stiff microgels* **(b)** *+RGDs (n≥14 per condition)* **(c)** *-RGDs (n≥11 per condition) (*p<0.05 by Kruskal-Wallis ANOVA). The box plots show 25/75th percentiles and the whiskers show the maximum and minimum data points. All data shown are mean± s.e.m*

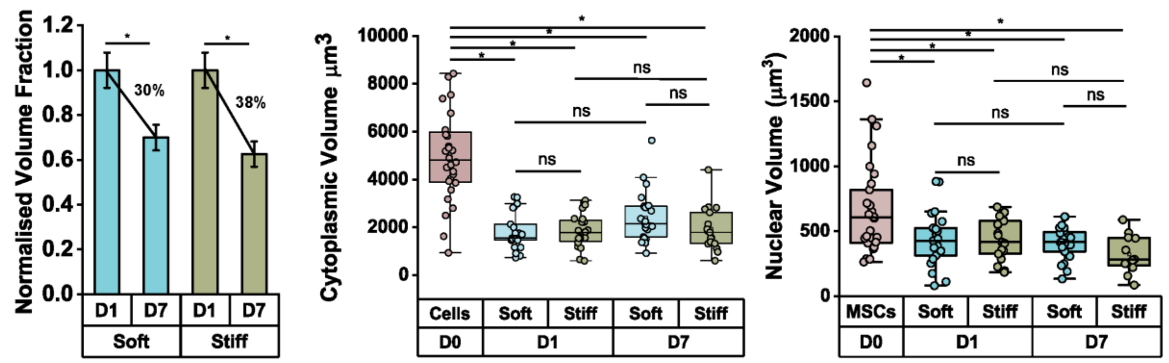

***Figure S4: (a)*** *Normalized volume fraction of MSCs in soft and stiff microgels over time* ***(b)*** Cytoskeletal volume ***(c)*** nuclear volume quantification of cells in soft and stiff microgels during proliferation for 7 days (n≥17) (*p<0.05 by Kruskal-Wallis ANOVA) The box plots show 25/75th percentiles and whiskers show the maximum and minimum data point.

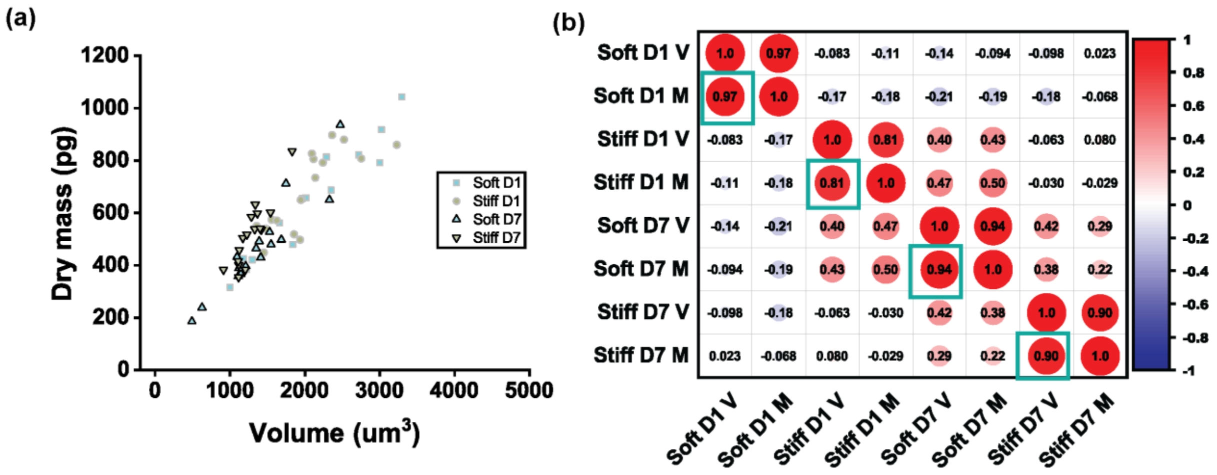


***Figure S5: (a)*** *Scatter plot of dry mass and volume of single cells in soft and stiff microgels.* ***(b)****. Pearson correlation plots of volume and dry mass correlation for soft and stiff microgels from D1 to D7. The (r) value (inside the circles) denotes how much is the correlation between two variables. A (r) value close 1 is a strong correlation (positive/negative) and close to 0 is a weak correlation to 1. The larger the circle is the more is the correlation. The dry mass and volume are positively correlated with values close to 1 which denotes they are highly dependent on each other. The green boxes denote the various time points compared between soft and stiff microgels (n≥15 microgels per condition)*

.

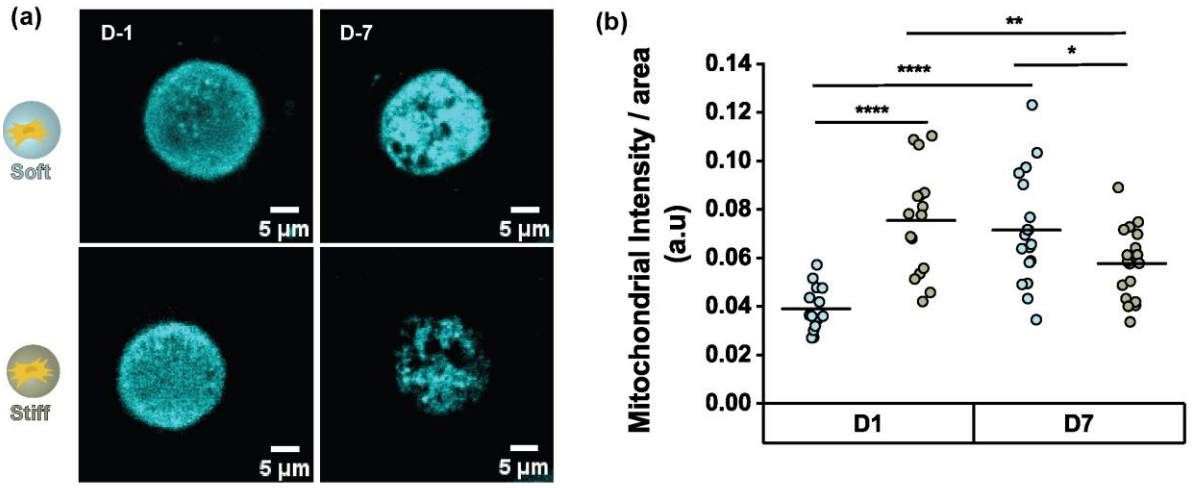

***Figure S6:*** *(a) Confocal micrographs and (b) quantification of mitochondrial concentration expression stained with Mitotracker (cyan) on cells in soft and stiff microgels up to 7 days (n≥15) (**** p<0.0001, **p<0.01, *p<0.5 by Kruskal- Wallis ANOVA).*

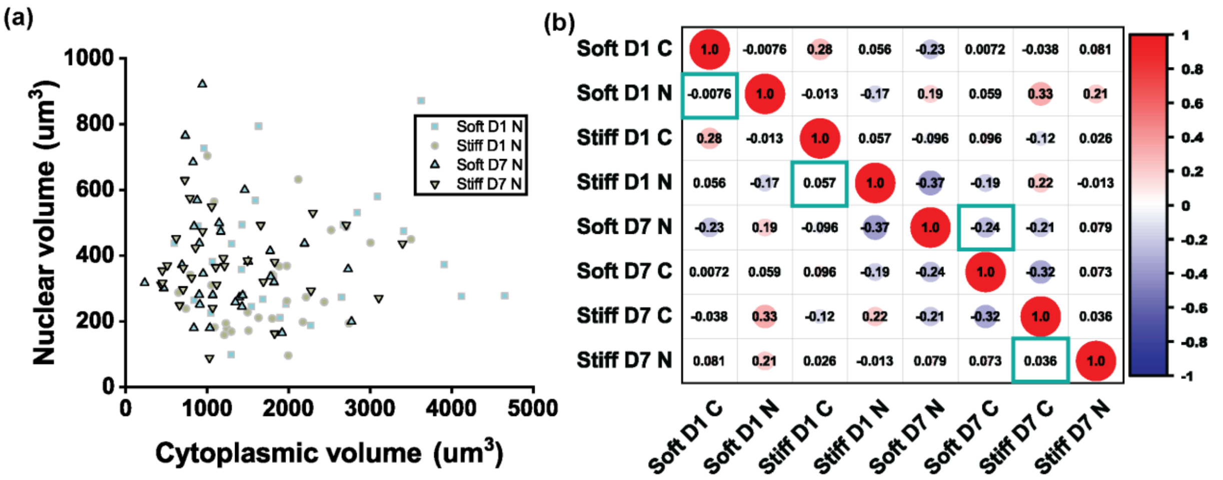


***Figure S7: a)*** *Scatter plot of dry mass and volume of single cells in soft and stiff microgels.* ***(b)****. Pearson correlation plots of volume and dry mass correlation for soft and stiff microgels from D1 to D7. The (r) value (inside the circles) denotes how much is the correlation between two variables. A (r) value close 1 is a strong correlation (positive/negative) and close to 0 is a weak correlation to 1. The cytoplasmic and nuclear volume are interestingly inversely between soft and stiff microgels. The soft microgels have a negative correlation between the cytoplasmic and nuclear volume while the stiff microgels have a positive correlation. The green boxes denote the various time points compared between soft and stiff microgels (n≥30 microgels per condition).*

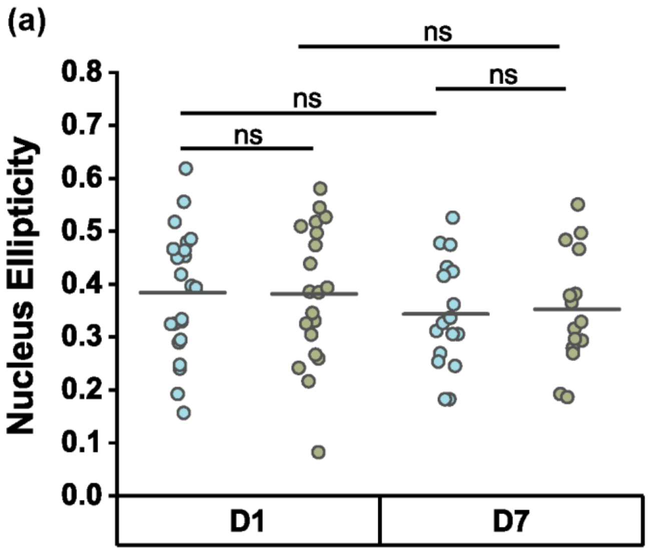


***Figure S8: (a)*** *Nucleus ellipticity obtained from 3D constructed micrographs up to 7 days from cells in soft and stiff microgels (n≥15 per condition).*
